# PAMR1 negatively impacts cell proliferation and migration of Human Colon Cancer HT29 Cell Line

**DOI:** 10.1101/2022.09.07.506931

**Authors:** Layla Haymour, Alain Chaunavel, Mona Diab Assaf, Abderrahman Maftah, Sébastien Legardinier

## Abstract

Colorectal cancer (CRC) is becoming one of the most prevalent cancers worldwide. Among cancers, it ranks the third place in terms of incidence and the second in terms of mortality. Even though immunological test allows fast and easy diagnostic method, there is no specific and reliable methods for early detection of CRC. Despite different treatments, high risk of re-occurrence is associated with advanced and metastatic CRC stages. An exhaustive knowledge on specific biomarkers or molecular actors involved in CRC could help to eradicate tumors or limit cancer recurrence. In this study, we focused on PAMR1 (Peptidase Domain Containing Associated with Muscle Regeneration 1), which is already considered as a tumor suppressor in breast and cervical cancers. *In silico* analysis of RNASeq data showed that PAMR1 was significantly downregulated in CRC tissues compared to their adjacent normal ones, as well as in cervical cancer. Our analysis showed that this downregulation, probably due to promoter hypermethylation, such as in breast cancer tissues, appeared in the four cancer stages as early as the first stage. In consistency with *in silico* analyses, the expression of PAMR1 was found to be lower at the transcript and protein levels in CRC tissue samples compared to normal ones, as well as in different CRC cell lines (HCT116, HT29, and SW620) compared to normal colon cell line (CCD841CoN). To understand the role of PAMR1 in CRC cancer, recombinant purified PAMR1 or concentrated secretome from CHO overexpressing PAMR1 were used to exogenously treat CRC cell lines with a focus on HT-29 cells as well as Hela cervical cancer cell line known to be sensitive to PAMR1. Transient or stable transfections were also performed to determine the impact of PAMR1 overexpression in HT29 and/or HeLa cells. In this study, we finally showed that presence of PAMR1 could reduce both cell proliferation and cell migration with a positive correlation between these biological effects and PAMR1’s quantity. This implies that PAMR1 expresses anti-proliferative and anti-migrative effects in CRC. Further studies to be done in order to confirm the tumor suppressive role of PAMR1 in CRC.

## INTRODUCTION

Cancer is becoming the most leading cause of death worldwide. Cancer diagnosis and patients’ treatment were impacted negatively with the Coronavirus disease 2019 (Covid-19) pandemic in Europe (Neamţiu et al., 2022) besides other factors that enhance its prevalence worldwide. According to the latest Global Cancer Observatory Statistics in 2020 (Globocan 2020), colorectal cancer (CRC) was classified the third cancer in terms of its incidence (1,931,590 cases), after lung and breast cancers, as well as the second leading cause of cancer death (935,173 cases), after lung cancer. In Europe, 4,398,443 new CRC cases (out of 19292789 cancer cases) were estimated in 2020. Although the risk of developing CRC is more pronounced after the age of 50 years (Byrne, 2017), environmental factors (Diergaarde et al., 2007) and genetic hereditary factors, such as Familial Adenomatous Polyps (Jasperson et al., 2010) and APC gene mutation (Valle, 2014), can be also incriminated in its occurrence and development. Screening of CRC can be assayed by stool-based, imaging and endoscopic tests (Hadjipetrou et al., 2017); as well as detecting tumor biomarkers, Carcinoembryonic Antigen (CEA) or Carbohydrate Antigen 19-9 (CA19-9), that are more or less specific for CRC. Despite various treatment methods of this malignancy, especially surgery in early stages, high mortality rate is associated with more advanced/metastatic stages. Have more knowledge on different actors, such as oncogenes and tumor suppressor genes, involved in CRC is still a challenge. Especially the finding of an early specific biomarker of CRC is a crucial issue for its early diagnosis, more targeted treatment, and high survival rate.

Peptidase Domain Containing Associated with Muscle Regeneration 1 (PAMR1) is a multi-domain secreted glycoprotein, formed of five main domains: CUB domain (Complement C1r/C1s, Uegf, Bmp1), one EGF-like domain (Epidermal Growth factor – like domain), two SUSHI domains (SUSHI 1 and SUSHI 2), and a trypsin-like peptidase S1 domain. PAMR1 was first shown to be downregulated in Duchenne Muscular Dystrophy (DMD) (Nakayama et al., 2004), with no clear idea about its regenerative mechanism of action. It was also reported to be suppressed in some cancers including breast cancer (Lo et al., 2015), cervical cancer (Yang et al., 2021) and gynecologic cancer (Yu et al., 2021). Studies focusing on PAMR1 in breast cancer turned out to show that PAMR1 is downregulated by means of epigenetic silencing due to its promoter’s hypermethylation (Lo et al., 2015). Recovering PAMR1’s expression by an epigenetic drug was shown to inhibit DNA methylation such as 5-aza-2’-deoxycytidine, leading to diminishing the invasion and migration of breast cancer cells (Lo et al., 2015). In this last study, PAMR1 was considered for the first time as a tumor suppressor. This role in cancer was recently confirmed by Yang et al. showing that PAMR1’s knockdown promoted proliferation, migration, and invasion of cervical cancer cells such as HeLa and Me180 cells (Yang et al., 2021). Despite these findings on PAMR1’s biological roles, the mechanism of action, including proteins partners, of this secreted glycoprotein is still unknown in skeletal muscle cells as well as in cancer cells. However, the presence of CUB and EGF-like domains in PAMR1 suggests its involvement in protein-protein interactions with other secreted proteins or cell-surface membrane proteins. Indeed, EGF-like domains are small protein domains (30-40 residues) stabilized by three disulfide bonds (Wouters et al., 2005) and known to regulate protein interactions such as those between Notch receptors and their Jagged and Delta-like ligands (Rand et al., 1997). In addition, The secreted protein SCUBE2 (secreted Signal Peptidase CUB-EGF domain containing protein 2), which exerts a tumor suppressor activity in breast cancer (Cheng et al., 2009), has similarly to PAMR1 a CUB domain and a multi-repeat region composed of 9 EGF-like domains, both involved in its anti-tumor effect (Cheng et al., 2009), (Lin et al., 2013). In addition to this CUB/EGF-like domains combination, the presence of *O*-fucose, known to modulate NOTCH-ligands interactions (Okajima et al., 2003)(Luther and Haltiwanger, 2009), on mouse PAMR1 was recently demonstrated in our lab (Pennarubia et al., 2020). However, the contribution of *O*-fucose in the function of PAMR1 has not yet been determined.

In spite of the absence of a clear view of PAMR1’s mechanism of action in cancer, its effects on different signaling pathways related to cell proliferation and cell survival were investigated. Indeed, Yang et. al illustrated ability of PAMR1 to suppress MYC and mTORC1 signaling pathways in cervical cancer (Yang et al., 2021). However, the role of PAMR1 and its mechanism of action may be different depending on the type of cancer where PAMR1 exerts a tumor suppressor role.

Starting with preliminary *in silico* analysis showing a reduced amount of *PAMR1* transcripts in colorectal cancer tissues from patients than in normal controls, we were interested to investigate whether PAMR1 exhibited a tumor suppressor activity in CRC such as in breast and cervical cancers. In addition of public data obtained by RNASeq methods, we confirmed PAMR1 down expression (qPCR, western blots) in tissue samples from CRC patients and in three different CRC cell lines (HCT116, HT29, SW620) *versus* a normal colon cell line (CCD841CoN). To study the potential role of PAMR1 in colorectal cancer, two main strategies were carried out, namely exogenous treatments of cancer cell lines with recombinant PAMR1 stably produced in mammalian CHO cells and transient (or stable) overexpression of the canonical isoform 1 of human PAMR1 in HT-29 cells as well as in cervical cancer HeLa cells. Since the production yield of human PAMR1 in stable CHO cells was too low for exogenous treatments, its murine counterpart exhibiting 90.3% identity with the mature human isoform 1 was used. To assess the relevance of the use of recombinant mouse PAMR1, cervical cancer HeLa cells were treated in parallel. The effects of PAMR1 treatment on cell viability, proliferation and migration were analyzed in both HT-29 and HeLa cancer cell lines.

## MATERIALS AND METHODS

### Database Analysis

RNA Seq data were extracted from the FireBrowse database (www.firebrowse.org), which examines various types of cancer by comparing tumor samples to normal ones. In this study, we focused on Colon Adenocarcinoma (COAD), Rectal Adenocarcinoma (READ), Colorectal Adenocarcinoma (COADREAD), and Cervical and Endocervical Cancers (CESC). PAMR1 expression levels were fused from COAD/READ/COADREAD or CESC.uncv2.mRNAs_normalized_log2.txt found in COAD or READ or COADREAD or CESC.mRNAs_Preprocess.level file. RNA Seq data were analyzed with PAST4 software.

### Clinical specimen

A panel of cancerous colorectal tissue samples with their corresponding cancer-adjacent tissues were retrieved from the archive of the CRB Limousin - CHU of Limoges. Ethics approval (CRB-CESSION-2021-008) was obtained from the “Comité médico-scientifique de la tumorothèque de l’Hôpital Dupuytren”, the bioethics committee of CHU of Limoges. Tissue samples used were classified either by colorectal cancer stages (I-IV) or by T classification of TNM staging. The clinicopathological information of the patients was also available.

### Cell Lines

Three human colorectal cancer cell lines were used in this study, namely HCT116 (ATCC CCL-247), HT29 (ATCC HTB38) and SW620 (ATCC CCL-227). Only one human cervical cancer cell line was also used, HeLa cells (ATCC CCL-2). HCT116, HT29, HeLa or derived-stable cell lines were cultured in DMEM growth medium (Gibco, Thermofisher Scientific). However, the SW620 cell line was grown in RPMI growth medium (Gibco, Thermofisher Scientific). The normal colorectal cell line CCD814CoN (ATCC CRL-1790), grown in EMEM growth medium (ATCC), was used with less than 15 passages. Flp-In^TM^ CHO cells (Thermo Fisher Scientific, Waltham, MA, USA) stably overexpressing mouse PAMR1, previously obtained (Pennarubia et al., 2020), were cultured in F12 growth medium. All culture media were supplied with 10 % Fetal Bovine Serum (FBS) (S1810 biowest, South America) and 0.5 % penicillin/streptomycin antibiotics (100 U/mL penicillin and 100 µg/mL streptomycin) (Gibco, USA). The cells were maintained at 37°C in a humidified atmosphere with 5 % CO2.

### Plasmid constructs

In order to produce the secreted forms of recombinant human PAMR1 (isoforms 1 & 2) in Flp-In^TM^ CHO- cells, we used the modified pSec-NtermHis6 vector containing secretory signal peptide (IgK Leader) as found originally in commercial vector pSecTag/FRT/V5- His-TOPO^R^ vector (Thermo Fisher Scientific, Waltham, MA, USA), but fused to six histidine residues (His6) and followed by Kpn I and BamH I cloning sites, as previously described (Pennarubia et al., 2018). This modified pSec-NtermHis6 vector was digested by Kpn I and BamH I and using the same strategy of prehybridized overlapping oligonucleotides as in the previous study (Pennarubia et al., 2018), a new cassette was inserted containing the sequence of V5 epitope, downstream of His6 tag, and new cloning sites Hind III and Xho I to generate new vector referred to as pSecPSHisV5. The cDNA sequences of human PAMR1 isoform 1 (NP_001001991.1) and isoform 2 (NM_001001991.3) without the signal peptides, were cloned between Hind III and Xho I restriction sites downstream of the sequence encoding N-terminal His6 and V5 tags. Resulting constructs named pSecPSHisV5-hPAM1 and pSecPSHisV5-hPAM2 harbored the sequence of human PAMR1 isoform 1 and human PAMR1 isoform 2, respectively. After nucleotide sequence verification, each plasmid construct was subjected to a cotransfection with pOG44 vector (Thermo Fisher Scientific, Waltham, MA, USA) expressing the Flp recombinase to produce stably transfected Flp-In^TM^ CHO cells (Thermo Fisher Scientific, Waltham, MA, USA). In order to overexpress human PAMR1 isoform 1 in HT29 cells, the commercial pcDNA3.1(+) (Thermo Fisher Scientific, Waltham, MA, USA) was used. The cDNA sequence of human PAMR1 isoform 1 (NP_001001991.1) was amplified by PCR from HEK total cDNAs and inserted downstream of the cytomegalovirus promoter of the vector using Hind III and Xho I cloning sites. The obtained recombinant vector was named pcDNA3.1-hPAMR1. The nucleotide sequence was verified before cells transfection.

### Cell culture and transfection

Recombinant human HisV5-PAMR1 isoforms 1 and 2 were produced by stable transfection of Flp-In CHO cells. Flp-In CHO cells were co-transfected with 1 µg of either pSecPSHisV5-hPAM1 or pSecPSHisV5-hPAM2 construct and 4 µL of the transfectant X-tremeGENE^™^ DNA Transfection Reagent (Sigma-Aldrich, Saint Louis, MO, USA) according to the manufacturer’s protocol. The selection started 24h post-transfection by Hygromycin B (Thermofisher Scientific, Waltham, MA, USA) of final concentration 500 µg/mL in F-12 medium. The recombinant PAMR1 produced by hygromycin-resistant cells was assessed by Western blot. Stable HT-29 cells overexpressing untagged human PAMR1 isoform 1 as well as the Mock cells were co-transfected with pcDNA3.1-hPAMR1 construct and empty pcDNA3.1 vector pcDNA3.1(+) (Thermo Fisher Scientific, Waltham, MA, USA) respectively, with transfectant X-tremeGENE^™^ DNA Transfection Reagent (Sigma-Aldrich, Saint Louis, MO, USA) according to the manufacturer’s protocol. The selection of transfected cells started 24h post transfection by changing DMEM growth medium containing 750 µg/mL of Geneticin (G-418). Different clones of Geneticin-resistant pool of cells overexpressing PAMR1 were selected and amplified. The level of PAMR1 expression in the Pool, Clones and Mock cells was assessed by qPCR.

### Real-Time quantitative PCR (qPCR)

Total RNA from tissue samples (after being grinded in Liquid Nitrogen) and cell lines was extracted using RNeasy mini kit (QIAGEN) and reverse transcribed using High-Capacity cDNA Reverse Transcription Kit (Thermofisher Scientific) according to the manufacturers’ protocol. qPCR was performed using TaqMan gene expression Master Mix (Thermofisher Scientific, Lithuania).

### Protein production and purification

Recombinant PAMR1 protein was produced from stably transfected CHO cells. Post cell seeding by 24h, the cells with >80 % confluency were washed with PBS 1X and cultured in fresh warm F-12 medium supplemented with 10 % FBS for 96h (optimal protein production with least degradation). The supernatant was then collected and proteins were precipitated in ammonium sulfate to reach 50 % saturation at RT and then centrifuged at 10,000g RT for 15 min. The precipitated proteins were purified based on nickel affinity purification by AKTA Prime Plus automated purification system (GE Healthcare). The sample passed over a 1 mL Nickel-Sepharose (HisTrap HP) affinity column at a flow rate of 1 mL/min according to a pre-recorded purification program. Using Buffer B (25 mM Tris-HCl, 500 mM NaCl, 500 mM Imidazole, pH 7.5), the sample was eluted by different imidazole concentrations. Different eluted fractions were established. The fractions that correspond to the peaks were tested by Coomassie blue staining and Western blot. The purest fractions containing PAMR1 were selected and concentrated in 10K Amicon (Sigma Aldrich, Ireland) by repeated centrifugation steps of 4500 G each at 4°C during 45 min each.

### Secretome concentration

Stable CHO-mPAMR1 cells that were already produced in the lab (Pennarubia et al., 2018), were seeded in 20 cm^2^ Petri dishes to become confluent after 24h. The culture media were aspirated and cells were washed twice with PBS 1X before adding 1 % FBS F-12 medium. After 48 h of incubation at 37°C, the supernatant was collected and centrifuged at 2500 g for 5 minutes to discard any floating dead cells. The supernatant was then concentrated 20 fold using Amicon 3K using several rounds of centrifugation, each was done at 4500 g, 4°C, for 45 minutes.

### Protein extraction and Western blot

Tissue samples were grinded in liquid nitrogen. Lysis buffer named RIPA (50 mM Tris-HCl, 150 mM NaCl, 1 % Triton X-100 (v/v), 0.5 % sodium deoxycholate (w/v), 0.1 % sodium dodecyl sulfate (v/v), pH 8) containing a cocktail of protease and phosphatase inhibitors (Roche Applied Science, Mannheim, Germany) was used to extract total proteins from tissue samples and CRC cell lines by its incubation with cell pellets for 1 h at 4°C. The lysates were centrifuged at 14,000 g for 15 min at 4°C. The concentration of proteins, in the supernatant, was quantified using BCA protein Assay Kit (Thermofisher scientific, USA). Proteins were resolved by SDS-PAGE in 8 % polyacrylamide gel at 24 mA. Then, proteins were blotted on 0.45 µm nitrocellulose membrane for 2 h at 50 mA. The membranes were blocked by TBS-Tween20 0.1 % (50 mM Tris, 150 mM NaCl, pH 7.6, 0.1% Tween-20 (v/v)) supplemented in 5 % Bovine Serum Albumin (BSA) (Sigma Aldrich, USA) or 5 % half-fat milk for 1h at room temperature. The membranes were then incubated with sheep anti-PAMR1 antibody (AF6517, R&D systems) diluted at 1:1000 in TBST 0.1 % supplemented with 2.5 % BSA or with Goat anti-GAPDH antibody (AF5718, R&D systems) diluted at in TBST-0.1 % supplemented with 2.5 % milk or anti-V5 HRP (Thermofisher scientific) overnight at 4°C. After washing the membrane with TBST-0.1 % three times, the corresponding secondary antibodies were added at 1:1000 for 1 h at room temperature. The membranes were revealed after adding the chemiluminescent substrate using an Amersham Imager 600 device (GE Healthcare, Uppsala, Sweden).

### Cell Viability Assay

Colorectal cancer cell lines were seeded to a cell density of 100,000 cells per well in 96-well plates. The cells were incubated for 24h in humidified condition at 37°C and 5% CO2. Different concentrations of purified recombinant PAMR1, diluted in growth medium, were added to cells, which were then incubated for 24 h, 48 h and 72 h. At each time point, 10 µL of cell counting kit (CCK8) (WST-8 CCK8, ab228554, abcam) was added in each well. The results were revealed by spectrometer at a wavelength of 460 nm 1 h post incubation with CCK8.

### Cell proliferation Assay

Cell lines were seeded at seeding density 500,000 cells per well in six-well plates. After 24 h, the medium was changed by adding different concentrations of purified PAMR1 or 20 folds concentrated stable CHO-mouse PAMR1 secretome. The cells were then incubated for 24 h, 48 h and 72 h. At each time point, cells were detached and added to the cells of the supernatant. Total cells were counted after being stained by Trypan blue using Malassez chambers.

### Cell Migration Assay

Cell migration assay was performed using two well-silicon inserts (Culture, Insert 2 well, Ibidi, Germany). Cells were put in the wells after being trypsinized and resuspended with growth medium supplemented with 10 % FBS. After the cells reached confluency, the inserts were removed, the cells were cultured in growth medium supplemented with 1% FBS. The closure of the cell-free gap was visualized daily and measured using Image J software.

### Statistical Analyses

All the experiments were performed independently at least three times. t-Student test found in GraphPad Prism 7 (GraphPad Software Inc, San Diego, CA, USA) was used to perform the statistical comparison. Results were considered as statistically significant if the p-value was less than 0.05.

## RESULTS AND DISCUSSION

### Low PAMR1 expression in colorectal cancer (CRC) and in many other cancers

Based on RNA Seq public data available in FireBrowse database (http://firebrowse.org/), *in silico* analysis comparing mRNA expression encoding many proteins in tumoral *versus* normal tissues can be done. For PAMR1, a down-expression was seen in colorectal cancer (COADREAD) and in many other cancers (Figure 1A), confirming previous findings for cervical (Yang et al., 2021), breast (Lo et al., 2015) and hepatocellular (Yin et al., 2016) cancers and also for cutaneous squamous cell carcinoma (Wei et al., 2018).

**Figure 1:**
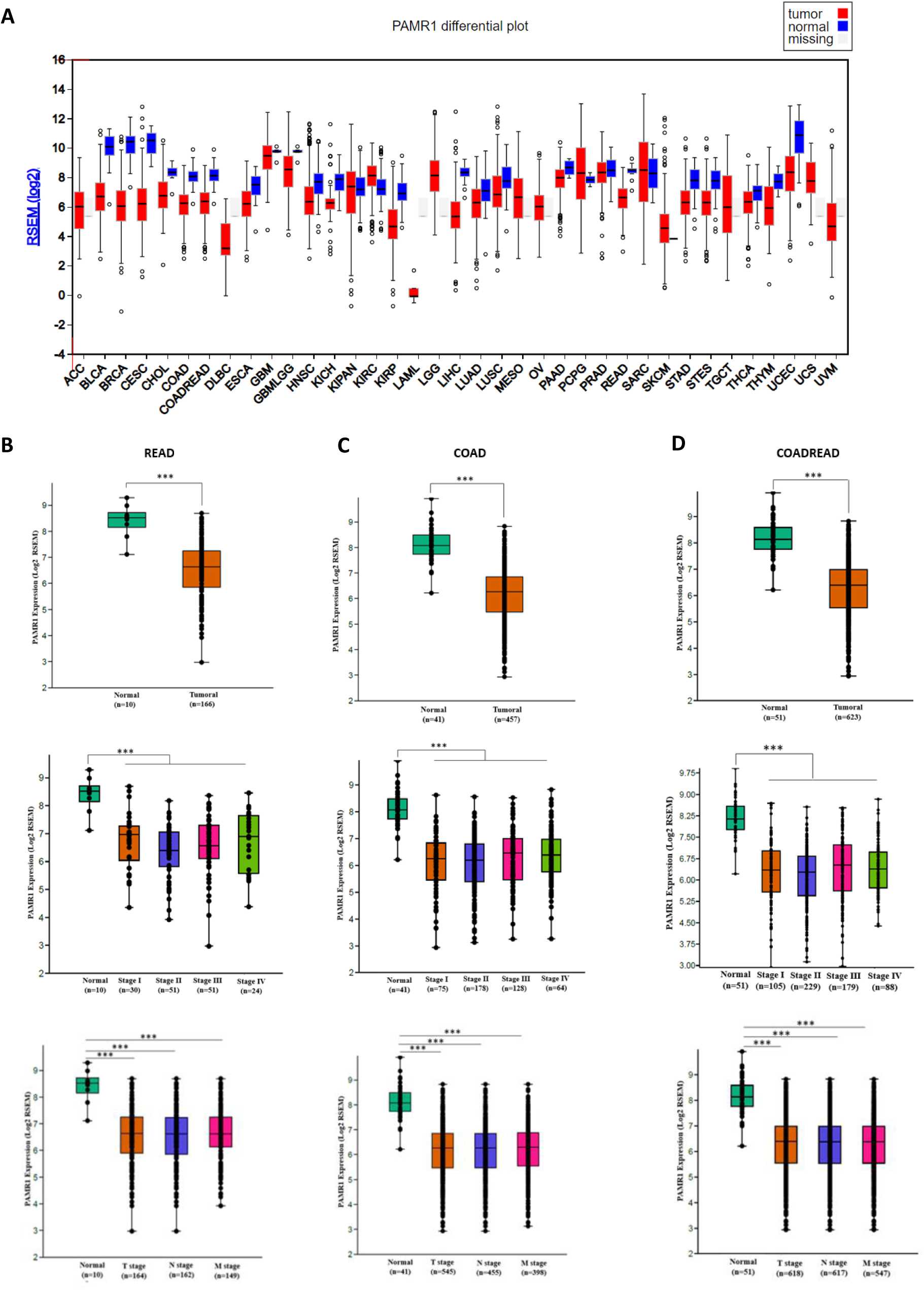
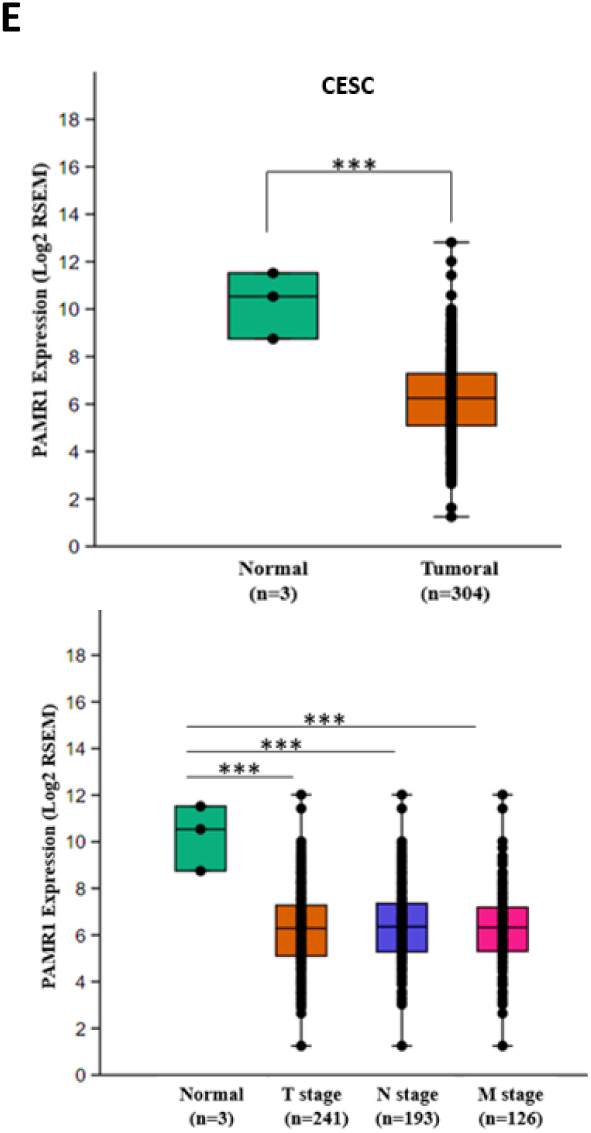
PAMR1 is suppressed in most cancers including colorectal and cervical cancers in stage-independent manner. (**A)** Expression of PAMR1 in normal and tumoral cancer samples from TCGA Firebrowse database (http://firebrowse.org/). PAMR1’s expression is downregulated in Cervical and Endocervical Cancers (CESC), Colon Adenocarcinoma (COAD), Colorectal Adenocarcinoma (COADREAD) and Rectal Adenocarcinoma (READ) tumoral samples compared to normal ones. (**B, C, D and E)** RNA Seq data Analysis showed a significant PAMR1 mRNA down expression in tumoral samples compared to normal samples in rectal, colon, colorectal adenocarcinomas and cervical cancer respectively. This downregulation is pronounced in all pathological stages (Stage I, Stage II, Stage III, Stage IV) and TNM stages, as early as stage I and T stage. The box plots represent the mean of Log2 RSEM ± SEM. The RNA Seq database analysis was carried out by PAST software. *p < 0.05, **p < 0.01, ***p < 0.001.

In this study, we especially focused on rectal adenocarcinoma (READ) (normal tissues =10 samples vs tumoral tissues = 166 samples) (Figure 1B) and colon adenocarcinoma (COAD) (normal tissues = 41 samples vs tumoral tissues = 457 samples) (Figure 1C) or on all compiled data available for colorectal adenocarcinoma (COADREAD) (normal tissues = 51 samples vs tumoral tissues = 623 samples) (Figure 1D). We were also interested in cervical cancer (CESC) (normal tissues = 3 samples vs tumoral tissues = 304 samples) (Figure 1E) exhibiting low quantity of PAMR1 and for which a tumor suppressor role was recently suggested (Yang et al., 2021). PAMR1 mRNA quantity was dramatically and significantly reduced in all these CRC tumoral tissues compared to their adjacent normal ones. Interestingly, this downregulation arose as early as stage I (or T stage according to TNM staging) and this low amount of PAMR1 mRNA was found in all the other stages analyzed. The stage-independent decrease of PAMR1 expression in cancer raises a question if PAMR1 could be considered as an early biomarker of colorectal cancer, such as in cervical cancer (Yang et al., 2021).

Strikingly, PAMR1 could be overexpressed in a few tumoral tissues such as the case of KIRC (Kidney renal clear cell carcinoma) and PCPG (Pheochromocytoma and Paraganglioma). However, due to missing data on normal tissues, we unfortunately could not evaluate the expression level of PAMR1 in the following cancers: ACC (Adrenocortical Carcinoma), DLBC (Lymphoid Neoplasm Diffuse Large B-cell Lymphoma), LAML (Acute Myeloid Leukemia), LGG (Brain Lower Grade Glioma), MESO (Mesothelioma), OV (Ovarian Serous Cystadenocarcinoma), TGCT (Testicular Germ cell Tumors), UCS (Uterine Carcinosarcoma), and UVM (Uveal Melanoma).

These very important differences in the expression of PAMR1 according to the type of cancer suggest that the role of this protein could differ according the cell type.

### Reduced expression of PAMR1 in CRC tissue samples from patients *versus* normal ones

In collaboration with CRB Limousin – CHU of Limoges, the expression of PAMR1 was analyzed at the transcript and protein levels in a panel of specimen collected from CRC patients, classified by pathological stages (Figure 2). According to qPCR results, PAMR1 expression was dramatically reduced in tumoral tissues from CRC patients compared to their adjacent non-cancerous ones (Figure 2A), confirming RNA seq data obtained from the FireBrowse database as shown in Figure 1. Only for the stage T1 of tumoral samples, a non-significant downward trend at transcript level was seen. Total proteins were also extracted from normal and CRC tissue samples followed by Western Blot using the same anti-PAMR1 antibody as in a previous study (Lo et al., 2015). The global analysis of three different sets of tissue samples showed a downward trend of PAMR1 quantity in tumoral samples for the four stages (Stages I-IV), as early as stage I (Figure 2B). However, PAMR1 was very difficult to quantify at the protein level, due to its low expression (even in healthy tissues), its instability and its propensity for degradation.

**Figure 2:**
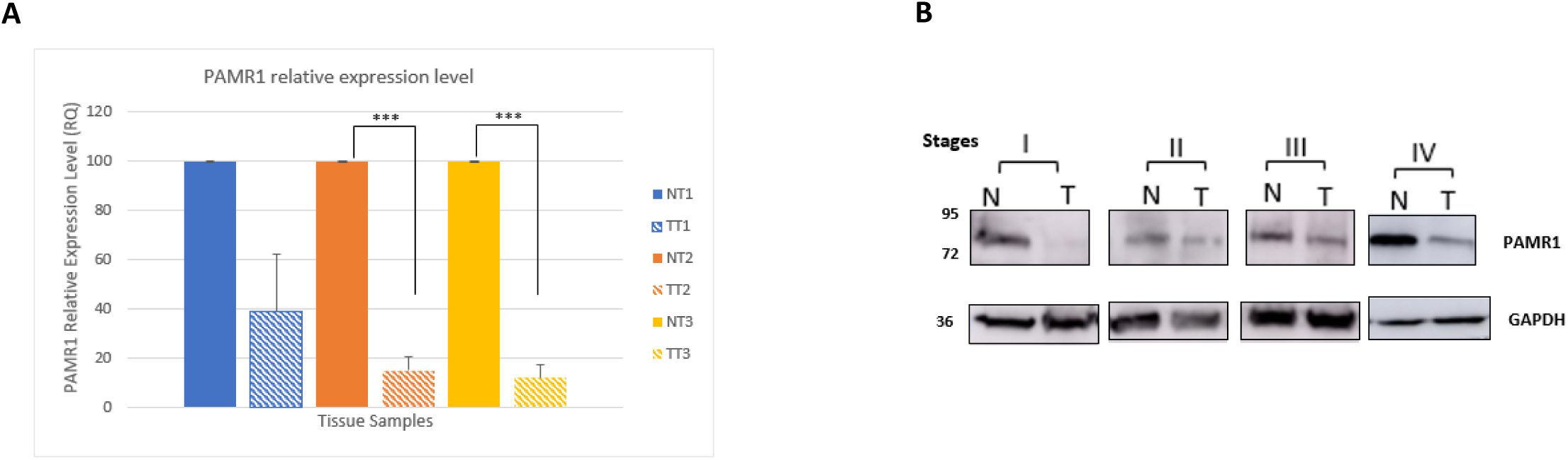
PAMR1 expression in CRC tissue specimen. (**A**) Bar graph showing PAMR1 relative expression level (ratio ± SEM for PAMR1/HSPA8) at transcriptomic level of three different sets of tissue samples classified by T stages (T1, T2, T3) of TNM classification. (**B**) Western blot analysis for PAMR1 expression in tumoral tissues from patients compared to their adjacent normal ones in the four CRC pathological stages. N: Normal. T: Tumoral. *p < 0.05, **p < 0.01, ***p < 0.001.

To conclude, all of these results showed at least a downward trend of PAMR1 expression in colorectal cancer at all stages. PAMR1 might be considered as an early biomarker of colorectal cancer as in cervical and breast cancers. This finding could help to limit the number of CRC patients diagnosed with aggressive stages (III and IV), correlated with poor prognosis and short overall survival time (Kuo et al., 2003)(Mukai et al., 2018).

### PAMR1 expression was significantly reduced in different colorectal cancer cell lines

The *in vitro* study was mainly based on the use of three colorectal cancer cell lines available in the lab, namely HCT116, HT-29 and SW620 cells, which represent colorectal cancer pathological stages I, II, and III, respectively. According to old Duke’s staging system, they are classified as Duke’s A, B, and C stages (Dukes, 1932) (Akkoca et al., 2014), respectively. Normal colon cell line CCD841CoN was used as normal control cells. In addition of these CRC cell lines, HeLa cells, representing the cervical cancer, were chosen in our study due to recent findings showing their sensitivity to a dysregulation of PAMR1 expression (Yang et al., 2021).

After extraction of total RNA from all the cell lines mentioned above, PAMR1 transcripts were specifically quantified by RT-qPCR using Taqman technology. As shown in Figure 3, the quantity of PAMR1 transcripts was found to be extremely lower (CT values above 35) in the three colorectal cell lines than in normal colon cell line CCD841CoN. If considering these high CT values, we can assume that PAMR1 expression was totally abolished in the three CRC cell lines as it was the case for HeLa cells studied here, consistent with the previous study on cervical cancer (Yang et al., 2021). The failure of detection of PAMR1 protein signal by Western blot from the crude secretome or intracellular proteins of these cell lines (data not shown) confirms qPCR results. Surprisingly, we were also not able to detect PAMR1 in crude secretome of CCD841CoN, probably due to low sensitivity of the antibody used or a too low secretion of PAMR1 in spite of its good expression at the transcript level.

**Figure 3:**
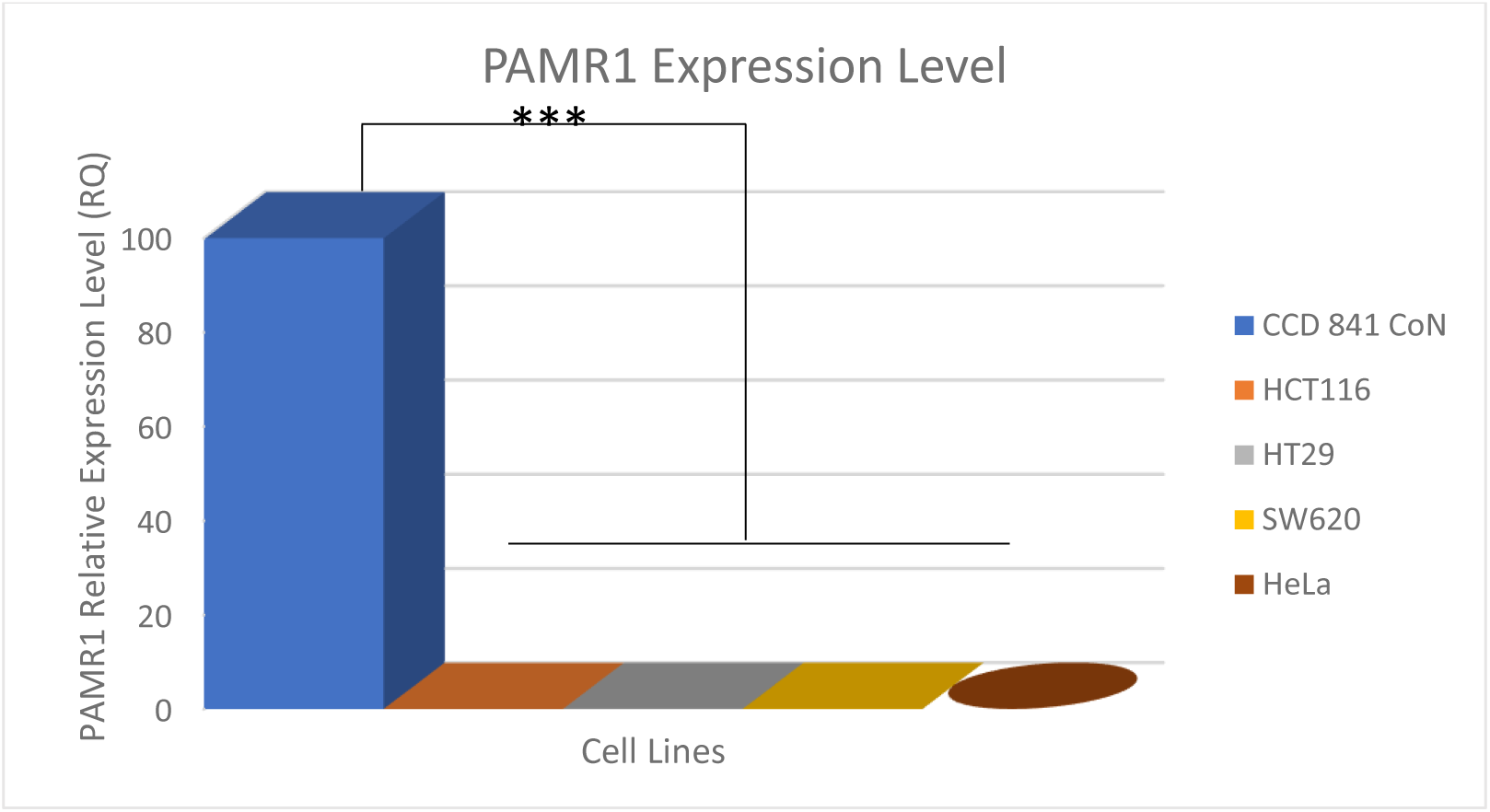
PAMR1 expression level in colorectal and cervical cell lines. PAMR1 transcripts are significantly downregulated in colorectal cancer cell lines (HCT116, HT29, and SW620), compared to normal colon CCD841CoN cells. Similarly, to CRC cell lines, cervical cancer HeLa cells also exhibited very low PAMR1 expression with high CT values around 36 (normal cervix cell lines not available). The histogram represents mean ± SEM. *p < 0.05, **p < 0.01, ***p < 0.001.

In compatibility with data from FireBrowse database, we thus ascertained the downregulated expression of PAMR1 in three colorectal cell lines of different stages and in HeLa cells, representing the cervical cancer.

### Production of recombinant PAMR1 for exogenous treatments of cancer cell lines

In a first approach, we wanted to produce recombinant human PAMR1 in order to carry out exogenous treatments of the three available CRC lines. These exogenous treatments required to produce human PAMR1 in an expression system and to purify it in the view of its addition to culture medium of non–modified cancer cell lines. As in our previous study (Pennarubia et al., 2020), stable CHO cell lines were generated to produce secreted forms of both isoforms 1 and 2 for human PAMR1, with N-terminal Histidine and V5 tags. Unfortunately, the expression level of both isoforms was much lower than for mouse HisV5-PAMR1, even undetectable for the canonical isoform 1 of human PAMR1 (data not shown). Thus, we chose the baculovirus insect cell system to express human PAMR1 isoforms 1 and 2. However, human PAMR1 expressed in the baculovirus-insect cell expression system was not very stable, prone to form aggregates and subjected to degradation both during its production and its purification. For all these reasons, only small quantities of the recombinant human PAMR1 were produced (data not shown) but did not allow to perform exogenous treatments of cancer cell lines with doses up to 5 µg/mL. Taking into consideration that mouse recombinant PAMR1 exhibits 90.3 % identity with the isoform 1 of human PAMR1 and whose level rate of production in stable CHO cells was much better, as previously shown (Pennarubia et al., 2020), we chose to rely on mouse PAMR1 to perform exogenous treatments. As shown in Figure 4A, significant differences of production were not seen in complete or serum-free medium for recombinant mouse PAMR1, detected by Anti-V5-HRP antibody around 95 kDa. However, a specific signal also appeared at about 50 kDa at 96h, probably due to partial protein degradation. Recombinant mouse PAMR1 was thus produced in complete medium and harvested after 96 h maximum followed by its purification on nickel affinity column (Ni-NTA). The analysis of the eluted fraction was done by Coomassie blue staining and Western blot (Figure 4B). By comparison to Western blot, the most enriched fractions in purified monomer of PAMR1 were fractions 9 and 10. Bands with high MW (up to 130 kDa) were seen and could correspond to protein aggregates that were formed either by interaction of PAMR1 with other protein partners or due to PAMR1-PAMR1 dimer/complex formation. These aggregates were not detected by the anti-V5 antibody, contrary to monomeric recombinant mouse PAMR1 which appeared at the expected size. The most enriched elution fractions in protein of interest (Figure 4B) were then pooled and concentrated before protein quantification using the BCA method.

**Figure 4:**
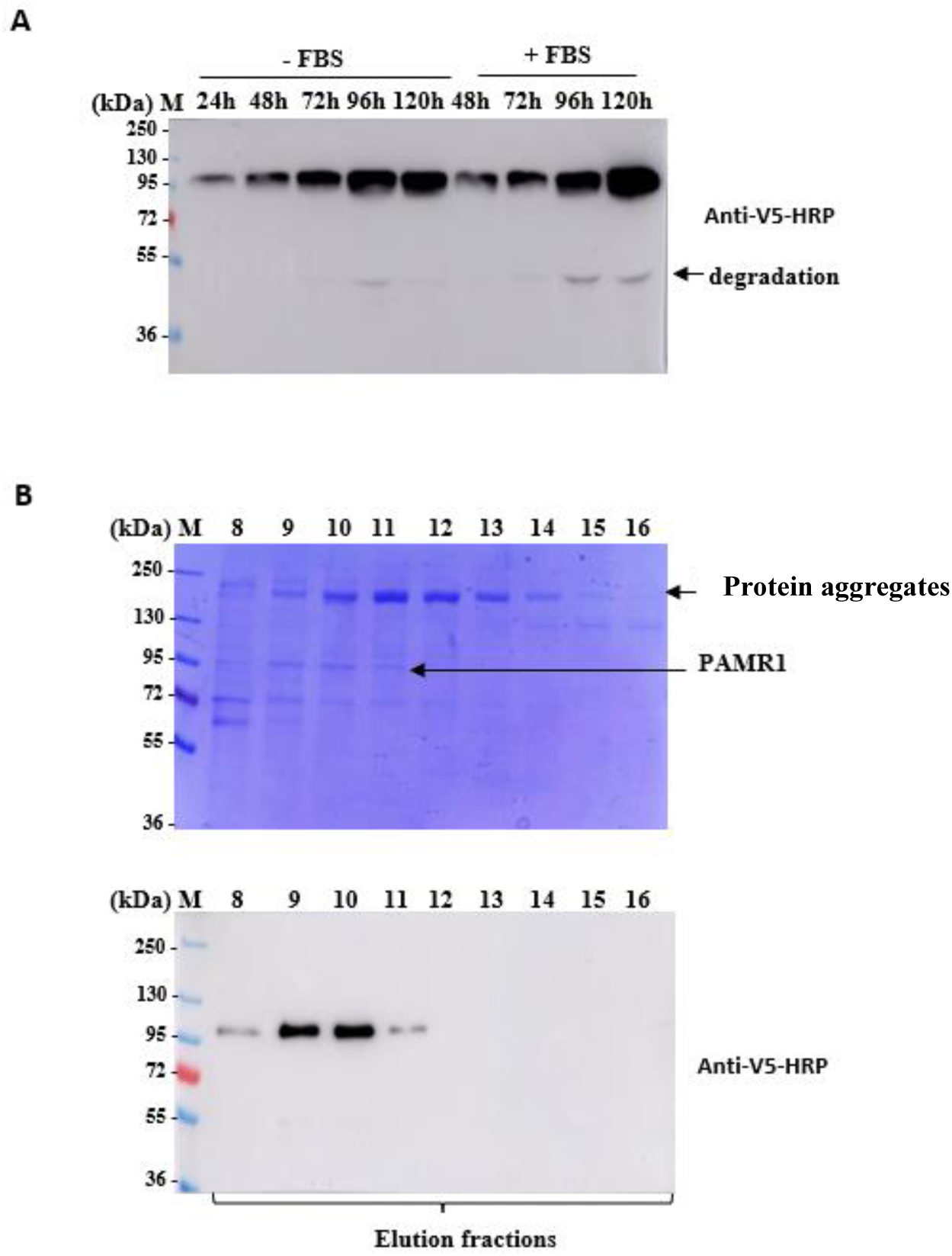
Purification of recombinant mouse PAMR1 produced by stable Flp-In^TM^ CHO cells. **(A**) Time course of production of recombinant mouse HisV5-PAMR1 in stable CHO cells in the presence and absence of FBS in Growth DMEM medium. **(B)** SDS-PAGE with Coomassie blue staining of polyacrylamide gel and Western blot analyses using anti-V5-HRP antibody of elution fractions following nickel-affinity purification using imidazole gradient of mouse HisV5-PAMR1, produced in secretome of stable CHO cells.

### HT-29 was the most sensitive cell line to exogenous treatment with recombinant PAMR1

Different concentrations of purified recombinant mouse PAMR1 (0, 1, 2.5, and 5 µg/mL) were added to culture medium of the three colorectal cancer cell lines, namely HCT116, HT29, and SW620 cells. To limit quantities of purified PAMR1 used, we chose to determine cell viability by using Cell Counting Assay 8 (CCK8) at different time points (0 h, 24 h, 48 h, and 72 h).

Exogenous treatment with purified mouse PAMR1 had no real effect on HCT116 and SW620 cells viability despite different doses and at different time points. However, the percentage of cell viability of HT29 decreased 48h and 72h post-treatment with the highest dose of PAMR1, namely 5 µg/mL. This suggests that PAMR1 either increased cell mortality and/or decreased cell proliferation. Compared to previous studies, it was tempting to think that PAMR1 exerted an anti-proliferative role with respect to HT29 (Figure 5). Cell viability assays were repeated several times, but with different preparations of purified mouse PAMR1. Each time, the results obtained ensure the sensitivity of HT29 to PAMR1. However, this effect was seen at different concentrations of purified recombinant PAMR1 depending on its purity in different preparations. This could be explained by the fact that, after concentration of elution fractions, purified mouse PAMR1 did not exhibit the same purity between preparations. Nevertheless, taking into account all of these results, we therefore chose to stably overexpress PAMR1 isoform 1 only in the HT-29 line, which exhibited a response to PAMR1 treatment.

**Figure 5:**
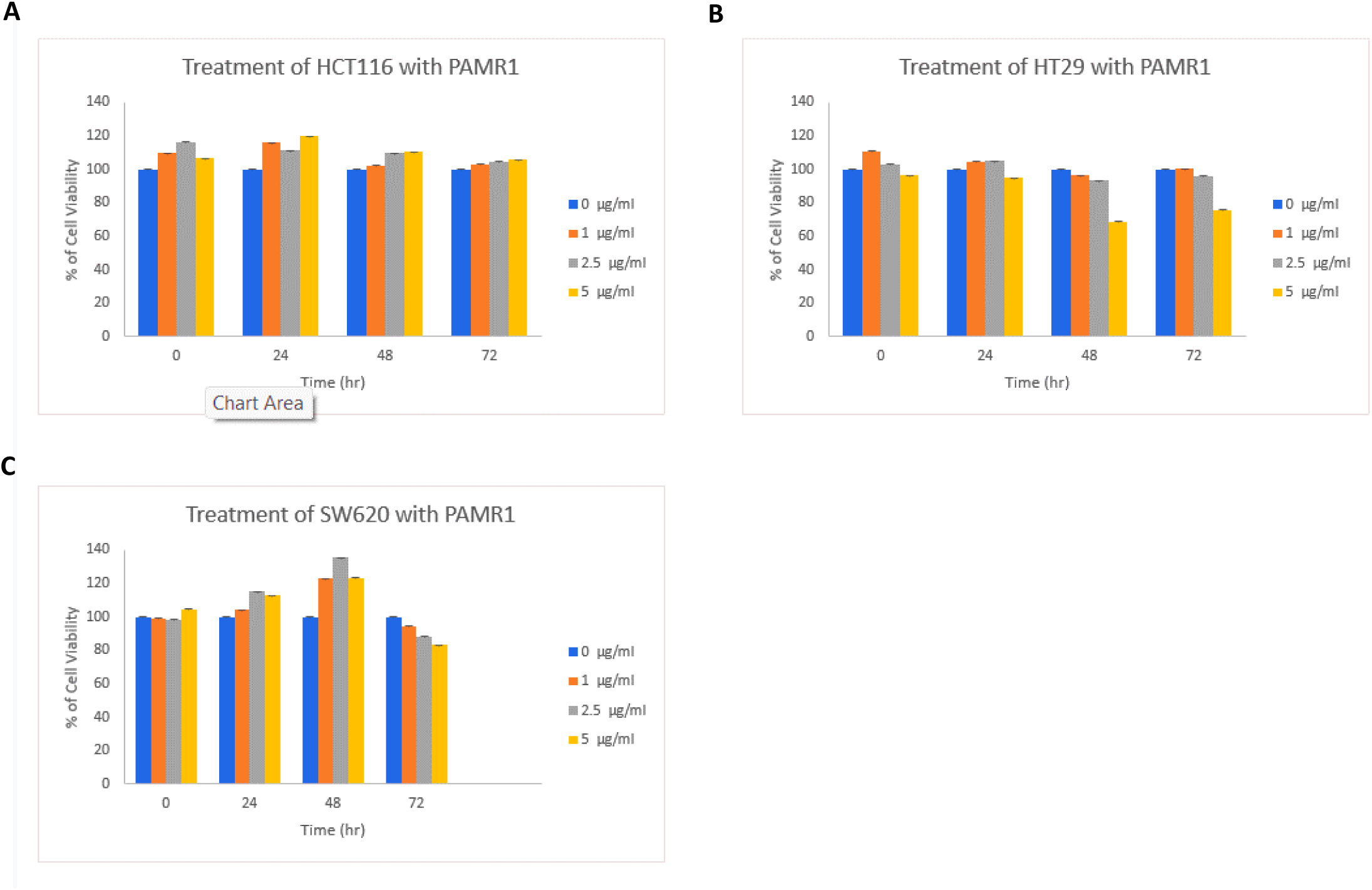
Cell viability assay (CCK8) for CRC cell lines. Representative histograms showing the percentage of cell viability ± SEM for HCT116 **(A)**, HT29 **(B)** and SW620 **(C)** cell lines, exogenously treated with different doses of purified recombinant PAMR1 (0, 1, 2.5 and 5 µg/ml), added at time point 0 h. Then, cells were incubated for 24, 48 or 72h before absorbance measurement of metabolized CCK8.

### Stable overexpression of human PAMR1 did not affect HT29 cell proliferation and migration

The priority of transfection went to HT29 that was found to be the most sensitive CRC cell line to exogenous treatment with recombinant PAMR1, compared to HCT116 and SW620. HT29 cells were first stably transfected with pCDNA3.1[hPAMR1 isoform1] or empty pCDNA3.1 vector. Hygromycin-resistant cells were selected and amplified to be analyzed by both RT-qPCR and Western blot (Figure 6). As expected, mock cells, represented hygromycin-resistant cells obtained after integration of empty vector, did not express PAMR1 at the transcript level (Figure 6A) and protein level using the anti-PAMR1 antibody (Figure 6B). However, the quantity of mRNA increased significantly for the pool and clone 2 (Figure 6A). The pool corresponded to a combination of several clones overexpressing different levels of PAMR1 whereas “Clone 2” from the pool was selected for its highest overexpression of human PAMR1 among all selected hygromycin-resistant clones. This reflects the overall low expression of PAMR1 in the Pool, with different clones exhibiting a relatively low expression level. The anti-PAMR1 antibody used for Western blot allowed the specific detection of 20 X concentrated overexpressed human PAMR1 isoform 1 at the expected size for the pool and clone 2 (Figure 6B). However, a strong specific band was also detected by anti-PAMR1 antibody at a higher apparent molecular weight around 120 kDa for clone 2. This suggests that beyond a certain concentration in the culture medium, PAMR1 was unstable and probably prone to form protein aggregates as seen for mouse PAMR1 purified after its ammonium sulfate precipitation (Figure 4B). This could also result from protein concentration of the secretome by ultrafiltration.

**Figure 6:**
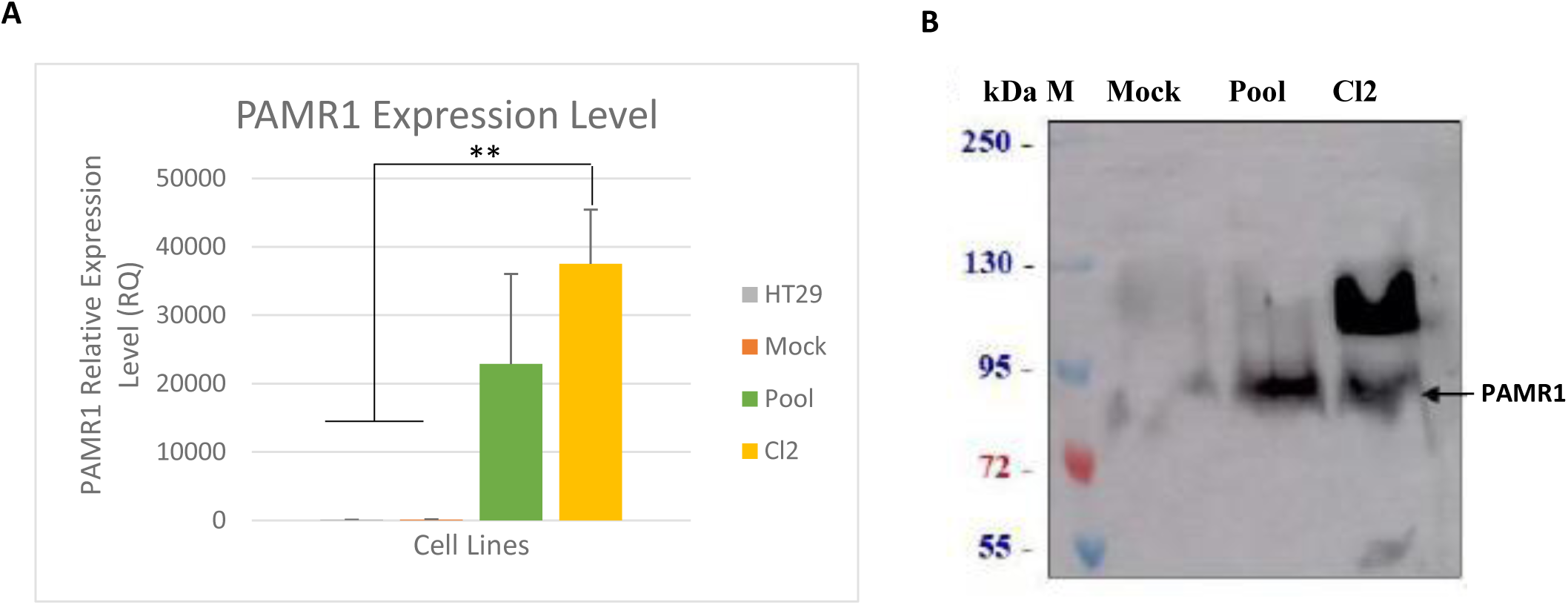
PAMR1 expression level in colorectal cancer HT29 cell line and derived stable cell lines. (A) PAMR1 expression at transcriptomic level in HT29 in comparison to stably transfected cell lines (Mock, Pool, and Cl2). The mRNA data of each cell line is normalized to its corresponding GAPDH mRNA level. The bar graph represents mean ± SEM. *p < 0.01, ** p < 0.05, ***p < 0.001. **(B)** Western blot analysis for PAMR1 protein expression in the concentrated secretome of Mock, Pool and Clone 2 (Cl2) using anti-PAMR1 antibody.

The biological effects of PAMR1 overexpression were assessed for all stably transfected cell lines and for non-modified HT29. No significant change was observed for cell proliferation (Figure 7A) and cell viability (Figure 7B) in stable cell lines overexpressing PAMR1 (Pool, clone 2) compared to WT HT29 or to Mock. For cell migration, the gap closure was almost seen at 144h for WT HT29 but not for the Cl2 and Pool stable cell lines, where the gap remained unclosed (Figure 7C). Thus, the quantification of gap closure showed a significant difference between stable cell lines and HT29 but not between cells overexpressing PAMR1 and Mock (Figure 7D). All these results could be explained by the insufficient quantity of PAMR1 expressed by stable cell lines compared to previous studies (Yang et al., 2021).

**Figure 7:**
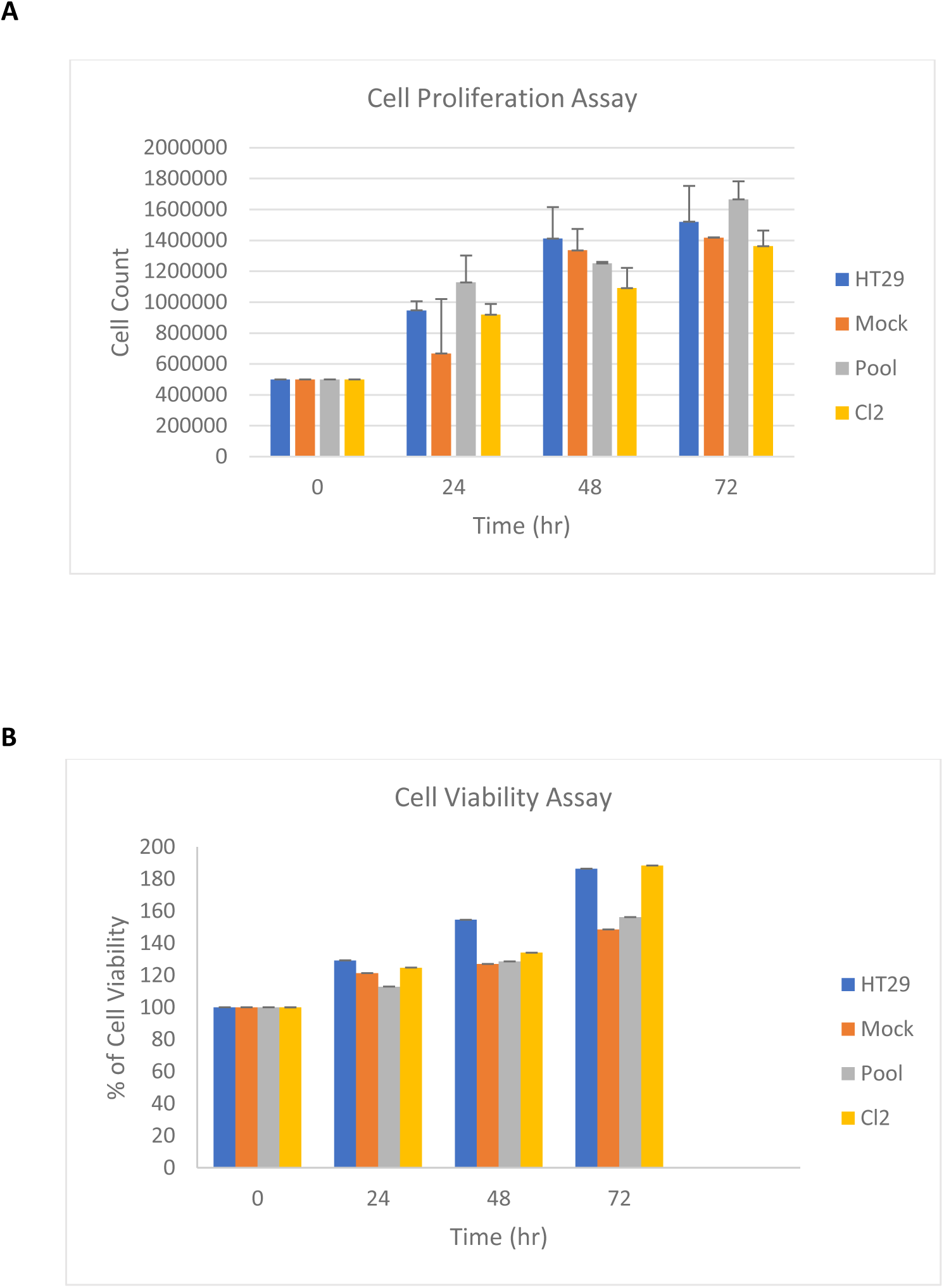
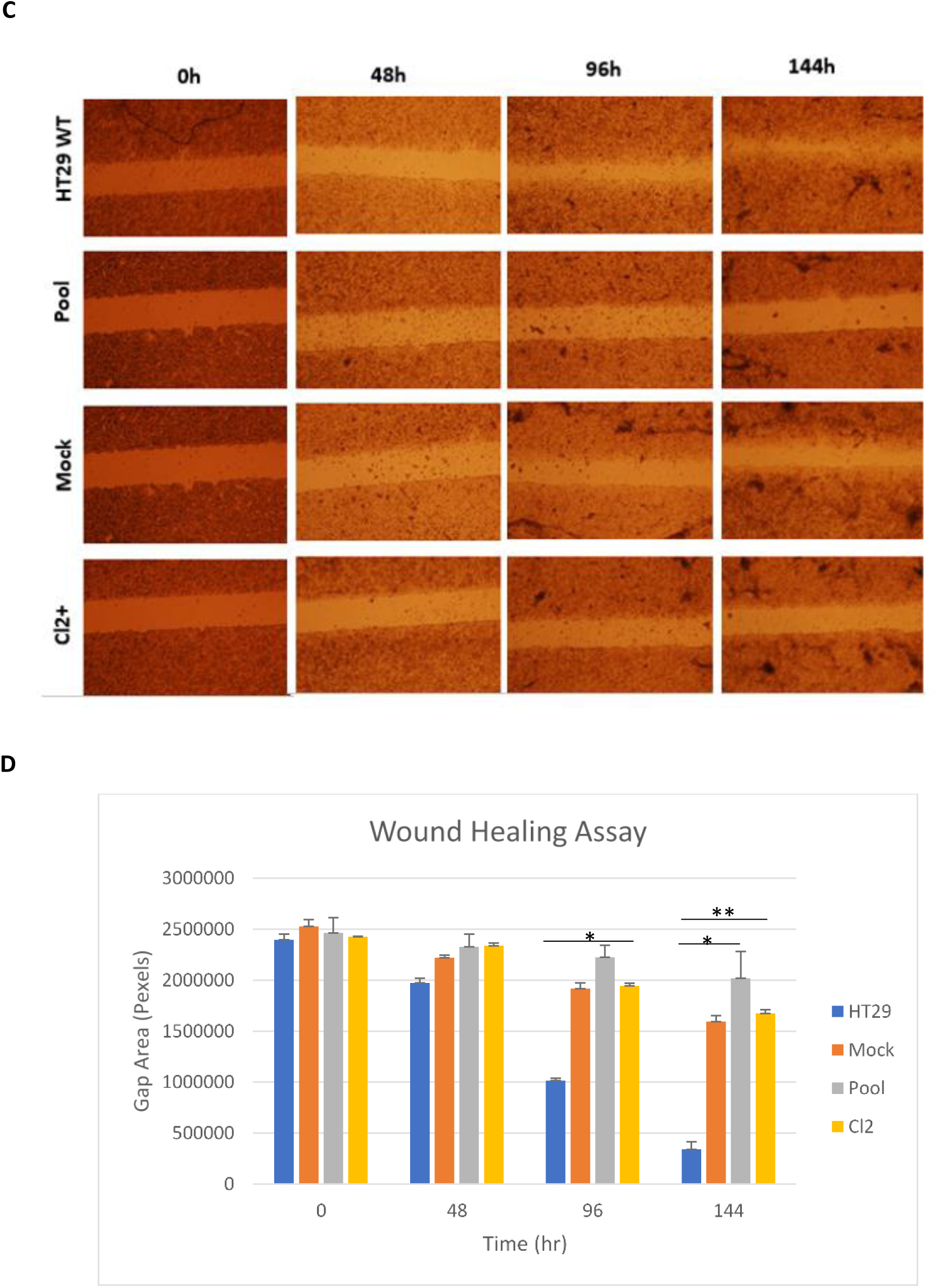
PAMR1 overexpression has no effect on cell viability, proliferation, and migration of stable HT29 hygromycin B-resistant cell lines. Cell proliferation **(A)** and cell viability **(B)** assays for HT29 stable cell lines (Mock, Pool, Cl2) compared to non-transfected HT29 cells, respectively. **(C)** Wound healing assays for HT29 and same stably transformed cell lines at different time points. **(D)** The bar graphs represent the mean of gap closure ± SEM. *p < 0.05, **p < 0.01, ***p < 0.001.

### The increase of PAMR1 quantity significantly reduced cell proliferation of HT29 cells

Since HT29 cell line was shown to be sensitive to recombinant mouse PAMR1 when added in culture medium but not to *in cellulo* human PAMR1 stable overexpression, we decided to combine transfection and treatment to increase the quantity of PAMR1 in the culture medium. Same experiments were done in parallel with HeLa cells, which are cervical cancer cell lines sensitive to PAMR1 dysregulation (Yang et al., 2021).

In a first approach, HT29 and HeLa cells were transiently transfected with the expression plasmid pCDNA3.1 bearing cDNA for human PAMR1 isoform 1 referred to as pCDNA-[hPAM iso1]. A second approach consisted on the overexpression of recombinant mouse PAMR1 in a stable CHO cell line already available in the lab (Pennarubia et al., 2020). The concentrated secretome of stable CHO cells was used instead of purified mouse PAMR1 to avoid protein degradation occurring during purification steps.

HeLa and HT29 were transiently transfected by pCDNA3.1-hPAMR1 iso1 construct, followed by detection of PAMR1 transcript by qPCR. The results showed a pronounced increase in PAMR1 expression at RNA level compared to non-transiently transfected cells with pCDNA3.1 (Figure 8). HT29 and HeLa cells were transiently transfected by pCDNA3.1[hPAM iso1] and/or either treated with concentrated recombinant PAMR1 (or with growth medium only). Then, cell proliferation was assessed after 72h of incubation by cell counting assay (Figure 9). The presence of PAMR1 whether being overexpressed by transient transfection or exogenous treatment provoked a significant decrease of cell number for HT29 (Figure 9A) but “slightly” reduced the cell proliferation of HeLa cells (Figure 9B). Consequently, the exogenous treatment of transfected cells by recombinant PAMR1 significantly reduced cells proliferation of both HeLa and HT29 cells. So, the combination of the two approaches highly increased the quantity of PAMR1, reflecting more efficient effect, thus more pronounced reduction of cell proliferation.

**Figure 8:**
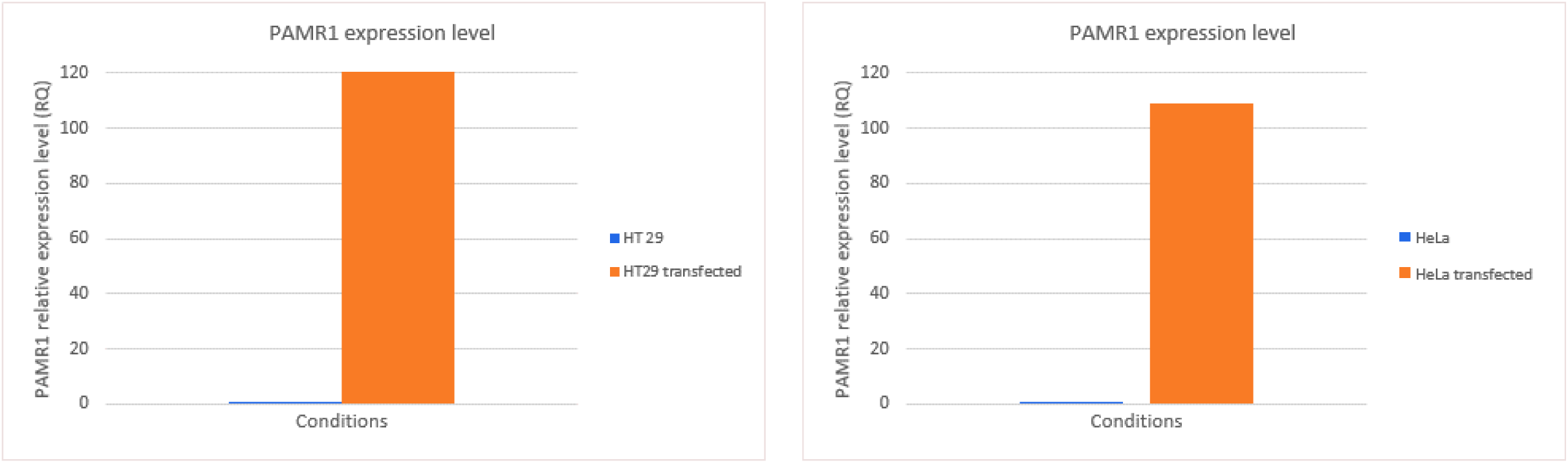
Overexpression of PAMR1 in transiently transfected cells. PAMR1 is overexpressed in transiently transfected cells by pCDNA- [hPAM iso 1] to a ratio 1:4 with respect to DNA transfection reagent. HT29 cells (left panel) and HeLa cells (right panel) compared to non-transfected ones.

**Figure 9:**
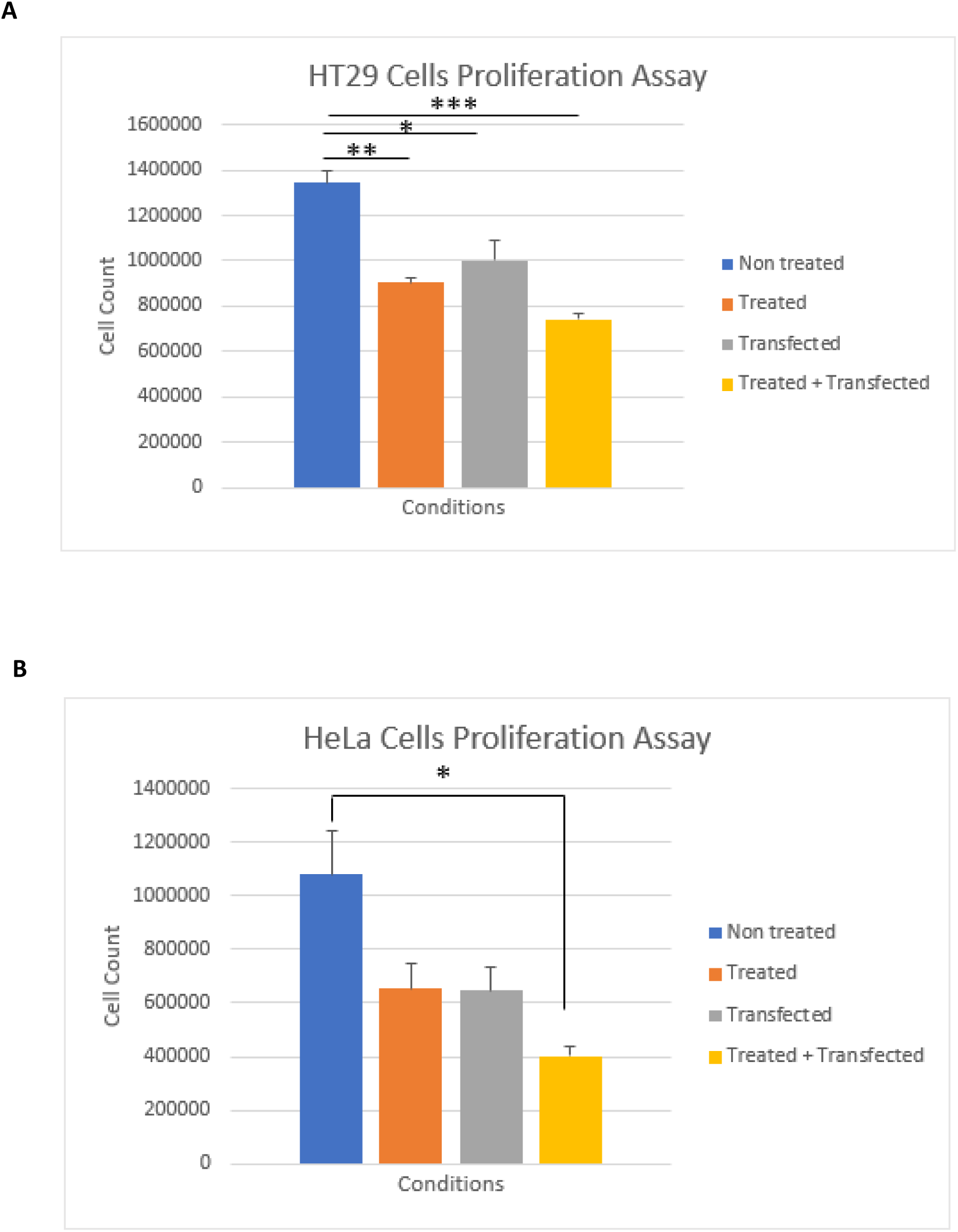
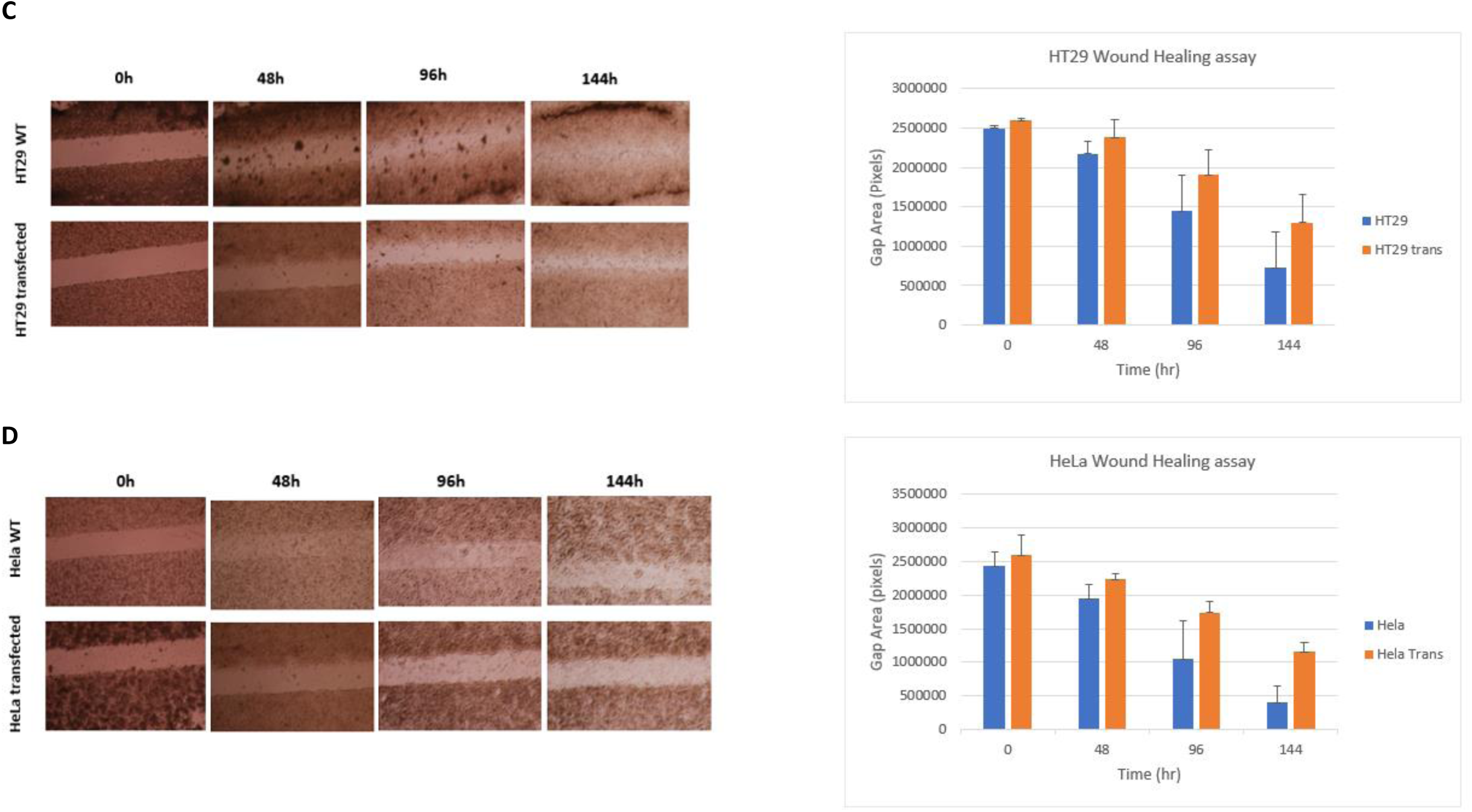
PAMR1 reduces HT29 and HeLa cells proliferation and migration. Cell proliferation assay for HT29 (A) and HeLa cells (B) in different conditions. Non treated: WT cells exogenously treated with concentrated CHO- mPAMR1 secretome. “Treated” corresponds to exogenously treated HT29 with concentrated CHO-mPAMR1 secretome. “Transfected” corresponds to transiently transfected cells by pCDNA3.1-hPAMR1. “Transfected + Treated” means that transfected cells were exogenously treated by concentrated CHO-mPAMR1 secretome. The bar graph represents mean ± SEM. * p < 0.05, **p < 0.001. Wound healing assays for non-transfected HT29 and transiently transfected HT29 (C) and HeLa (D) cells at different time points. The bar graphs represent the mean of gap closure ± SEM. *p < 0.05, **p < 0.01, ***p < 0.001.

Cell migration of transiently transfected HT29 and HeLa cells with their control (non-transfected cells) was assessed by wound-healing assay (Figures 9C and 9D). The gap closure started after 48h for non-transfected HeLa and HT29 cells and became more pronounced after 144h hours, whereas less cell migration was observed for the transfected ones. The results obtained took more than 48h to be visualized and were not significant reflecting that PAMR1 expression level is low and not sufficient to exert a pronounced effect within a short time duration.

To conclude, a significant increase of PAMR1 amount in the secretome of colorectal cancer cell lines, as well as in cervical cancer ones, significantly diminished cell proliferation but only a downward trend was observed for cell migration of both cell lines. These biological effects seem to be dependent of the level expression or quantity of PAMR1. This confirms the induction of HeLa and Me180 cells proliferation, migration and invasion as a result of PAMR1 knockdown. On the other hand, a huge PAMR1 overexpression in these cell lines reduced these effects.(Yang et al., 2021).

### Silencing PAMR1 expression in CRC might be due to promoter hypermethylation

Molecular events leading to PAMR1 downregulation in colorectal cancer as early as in stage I is unidentified yet; however, recovering its expression could be through the use of drug treatments as the case in other cancers (Lo et al., 2015) and with other suppressed genes (Zhu et al., 2018). Recovering the tumor suppressor effect of PAMR1 in breast cancer was through treatment with demethylation agent, 5-aza2’deoxycytine2 (Lo et al., 2015). Since PAMR1 was inactivated due to epigenetic silencing through promoter hypermethylation in breast cancer cells, 5-aza2’deoxycytine2 led to PAMR1 re-expression, thus reduction of cancer cell growth. To assess whether PAMR1 downexpression in CRC is due to its epigenetic silencing, treatments with demethylation agents could be performed. However, this treatment is not specific of PAMR1 and other tumor suppressor genes might be re-expressed following the use of this non-specific demethylating agent.

### The precise mechanism action of PAMR1 remains to be elucidated

The mechanism of action of PAMR1 is still unknown but PAMR1, which is a secreted multi-domain protein, could interact with one or several proteins expressed on the cell surface or present in the extracellular space. We can hypothesize that its protein partners, potentially different according to the cell type, could modulate its action. In the physiological state, PAMR1 could participate to the maintenance of a normal proliferation rate of different cell types. However, the suppression of its expression, by epigenetic inactivation or by other molecular events, in cancer cells undoubtedly participates to their increased proliferation. PAMR1 was thus recently considered as a tumor suppressor gene (Yang et al., 2021).

In conclusion, we confirmed the down expression of PAMR1 in colorectal cancer. The overexpression of PAMR1 is crucial for reduction of cell proliferation and migration of colorectal cancer cells. By that, PAMR1 could be predicted as an early biomarker and tumor suppressor of colorectal cancer. However, its mechanism of action is to be investigated.

## Acknowledgements

We acknowledge funding support from “Comité départemental de la Haute-Vienne, de la Creuse et de la Corrèze de la Ligue contre le cancer, France”.

